# Heparanase promotes Syndecan-1 expression to mediate fibrillar collagen and mammographic density in human breast tissue cultured *ex vivo*

**DOI:** 10.1101/2020.06.04.135202

**Authors:** Xuan Huang, Gina Reye, Konstantin I. Momot, Tony Blick, Thomas Lloyd, Wayne D. Tilley, Theresa E. Hickey, Cameron E. Snell, Rachel K. Okolicsanyi, Larisa M. Haupt, Vito Ferro, Erik W. Thompson, Honor J Hugo

## Abstract

Mammographic density (MD) is a strong and independent factor for breast cancer (BC) risk and is increasingly associated with BC progression. We have previously shown in mice that high MD, which is characterised by the preponderance of a fibrous stroma, facilitates BC xenograft growth and metastasis. This stroma is rich in extracellular matrix (ECM) factors, including heparan sulfate proteoglycans (HSPGs), such as the BC-associated syndecan-1 (SDC1). These proteoglycans tether growth factors, which are released by heparanase (HPSE). MD is positively associated with estrogen exposure and, in cell models, estrogen has been implicated in the upregulation of HPSE, the activity of which promotes SDC expression. Herein we describe a novel measurement approach (single-sided NMR) using a patient-derived explant (PDE) model of normal human (female) mammary tissue cultured *ex vivo* to investigate the role(s) of HPSE and SDC1 on MD. Relative HSPG gene and protein analyses determined in patient-paired high versus low MD tissues identified SDC1 and SDC4 as potential mediators of MD. Using the PDE model we demonstrate that HPSE promotes SDC1 rather than SDC4 expression and cleavage, leading to increased MD. In this model system, synstatin (SSTN), an SDC1 inhibitory peptide designed to decouple SDC1-ITGαvβ3 parallel collagen alignment, reduced the abundance of fibrillar collagen as assessed by picrosirius red viewed under polarised light, and reduced MD. Our results reveal a potential role for HPSE in maintaining MD via its direct regulation of SDC1, which in turn physically tethers collagen into aligned fibres characteristic of MD. We propose that inhibitors of HPSE and/or SDC1 may afford an opportunity to reduce MD in high BC risk individuals and reduce MD-associated BC progression in conjunction with established BC therapies.

## 1 Introduction

In Australia, mammographic screening reduces mortality by around 50% for screening participants (Roder, Houssami et al. 2008, Nickson, Mason et al. 2012). However, mammographic test sensitivity is impaired in women who have high mammographic density (HMD), where the dense area on the mammogram can mask BC-associated features, contributing to an increase in ‘interval cancers’ that arise within 1-2 years of a “clear” mammogram, and an increase in false negative and false positive screens (Nelson, O’Meara et al. 2016). HMD is therefore an important impediment to effective screening.

After adjusting for age and body mass index (BMI), on a population basis HMD is also one of the strongest risk factors for BC. BC is currently diagnosed in 1 in 8 Australian women over their lifetime, and accounts for approximately 19,500 cases and 3,000 deaths annually (AIHW 2019), making BC the major cause of female cancer-associated death in Australia. HMD is common; approximately 43% of women aged between 40 and 74 have heterogeneously or extremely dense breasts, wherein HMD regions represent greater than 50% of the total area (Sprague, Gangnon et al. 2014). An underappreciated fact is that HMD-associated BC risk is more impactful than other known risk factors, including the BRCA1/2 BC predisposition genes, when considered on a population-wide basis (Hopper 2015). In addition to increased BC risk, evidence is also emerging that HMD is associated with increased BC recurrence (Hwang, Miglioretti et al. 2007, Cil, Fishell et al. 2009, Shawky, Ricciardelli et al. 2015, Huang, Chen et al. 2016, Shawky, Huo et al. 2019) and treatment resistance (Elsamany, Alzahrani et al. 2015). Thus, interventions that reduce MD may offer new avenues for the prevention, diagnosis and treatment of BC.

A standout molecular feature of HMD is the preponderance of extracellular matrix (ECM), largely comprised of collagen fibres and proteoglycans (PGs), which are carbohydrate-coated proteins (Huo, Chew et al. 2014, Shawky, Ricciardelli et al. 2015). Abnormalities in ECM have been implicated in many pathologies, including cancer (Lu, Weaver et al. 2012). Importantly, targeting ECM components has proven efficacious for the treatment of some diseases (Jarvelainen, Sainio et al. 2009, Ferro, Liu et al. 2012). In the breast, increased abundance and organisation of collagen is associated with HMD (Boyd, Martin et al. 2010, Huo, Chew et al. 2015, McConnell, O’Connell et al. 2016) and our LMD vs HMD ‘within breast’ comparative data have confirmed that collagen-rich ECM is the most discriminatory feature of HMD (Lin, Cawson et al. 2011, Huo, Chew et al. 2015). Accumulation of collagen can influence mammary malignancy both *in vitro* (Provenzano, Inman et al. 2008, Levental, Yu et al. 2009) and *in vivo* (Provenzano, Eliceiri et al. 2009), and the collagen profile of HMD predicts poor survival in BC (Conklin, Eickhoff et al. 2011). Although dense collagen is undoubtedly a major feature of HMD-associated ECM, it forms a scaffold for a wide range of around 350 distinct ECM proteins, collectively called the ‘Matrisome’, which includes many PGs (Naba, Clauser et al. 2012). HMD stroma has many similarities with BC-associated stroma, where both promote the progression of malignancy. To date, the specific components of HMD ECM that promote BC are unknown. One candidate of interest, SDC1, has been positively associated with both HMD and BC (Shawky, Ricciardelli et al. 2015). HMD is approximately 60 % genetically inherited (Boyd, Dite et al. 2002), and GWAS studies have identified single nucleotide polymorphisms (SNPs) in several PGs and in PG-modifying enzymes that correlate with increased BC risk (Shawky, Ricciardelli et al. 2015). In particular, we have identified SDC1 and SDC4 SNPs in BC (Okolicsanyi, Buffiere et al. 2015).

The heparan sulfate proteoglycan (HSPG) family of glycoproteins include membrane bound proteins SDC1 and SDC4, glypican (GPC 1-6), beta-glycan, neuropilin-1 and CD44 (hyaluronic acid receptor), secreted extracellular matrix components (agrin, perlecan, type XVIII collagen), and the secretory vesicle molecule, serglycin (Sarrazin, Lamanna et al. 2011). HSPGs are widely involved in biological activities, including cell signalling, cell adhesion, ECM assembly, and growth factor storage (reviewed in (Nagarajan, Malvi et al. 2018)). HSPGs display primarily heparan sulfate-containing glycosaminoglycan (GAG) side chains that bind growth factors, which are in turn released by the action of HPSE (Shawky, Ricciardelli et al. 2015). This enzyme trims HS GAG chains from HSPGs (including SDC1), allowing the PG core proteins to be cleaved by enzymes such as MMP-9. For SDC1, this results in release from the cell membrane. Furthermore, a positive feedback loop occurs whereby SDC1 shedding stimulates its own expression (Ramani, Pruett et al. 2012). Shed SDC1 is also taken up by BC cells to promote proliferation (Su, Blaine et al. 2007), and has been shown to be important in Wnt signalling, contributing to tumourigenesis (Alexander, Reichsman et al. 2000).

HPSE expression strongly correlates with poor survival in BC (Sun, Zhang et al. 2017) and serum levels of shed SDC1 have been identified to be informative in regards to progression of several cancers including breast (Joensuu, Anttonen et al. 2002, Vassilakopoulos, Kyrtsonis et al. 2005, Szarvas, Reis et al. 2016). Shed SDC1 abundance detected in serum has been associated with BC size (Malek-Hosseini, Jelodar et al. 2017), and to be a result of chemotherapy (Ramani and Sanderson 2014). SDC1 gene expression in BC can be prognostic (Cui, Jing et al. 2017), with its induction in stromal fibroblasts identified in invasive BC (Yang, Mosher et al. 2011, Vlodavsky, Beckhove et al. 2012). Of relevance to BC, HPSE action to deglycanate SDC1 is essential for the binding of the core protein of SDC1 with lacritin, a protein known to be expressed in the breast, enabling mitogenic signaling (Weigelt, Bosma et al. 2003, Ma, Beck et al. 2006).

Recent data identifying a positive association of HMD with lifetime exposure to estrogens is also a key finding to our understanding of MD in BC (Huo, Chew et al. 2015). Estrogen unequivocally promotes MD, as evidenced by data showing alterations of MD with hormone contraceptives, hormone replacement therapy (Greendale, Reboussin et al. 1999, Byrne, Ursin et al. 2017), menopause (Stone, Warren et al. 2009), and treatment with anti-estrogenic drugs (Chew, Huo et al. 2014, Shawky, Martin et al. 2017). Estrogen also influences cells within the tumour microenvironment, causing immunosuppression (Rothenberger, Somasundaram et al. 2018). However, the mechanistic basis of estrogen promotion of MD is, as yet, unknown. Crucially, however, estrogen can directly upregulate HPSE in MCF-7 cells (Elkin, Cohen et al. 2003, Xu, Ding et al. 2007), and thus the E2 /HPSE /SDC1 axis has strong potential as the mediator of MD effects on BC through autocrine/paracrine pro-malignant actions.

We hypothesized that estrogen promotes HMD by up-regulating HPSE and SDC1 as part of its pro-oncogenic effects. Using anti-estrogens as well as HSPE inhibitors developed for cancer treatment and in use in clinical trials, and the SDC1 specific inhibitor SSTN, we investigated their effects in patient-derived explants of normal mammary tissue cultured over a two-week period. Our data identified that MD change was detectable and linked to the estrogen-HPSE-SDC1 axis.

## 2 Materials and Methods

### 2.1 Patient cohort

Human breast tissue specimens with no evidence of malignancy, and surplus to pathology needs, were accrued from prophylactic mastectomy surgeries, primarily in women with high BC risk, as determined by family history of breast cancer and/or with contralateral benign or malignant disease. Tissue from patients on recent (<6 months) hormone-based therapies or patients who carried BRCA1/2 mutations were excluded from the study. The demographics of the patients who donated their breast tissue for this study are detailed in Table 1. The study was approved by the Metro South Hospital and Health Services, Queensland (HREC/16/QPAH/107). Resected breast tissue was placed on ice and taken to the Pathology Department at the respective hospital, cut into 1-1.5 cm thick slices and checked for lesions. Up to 3 slices were then obtained and used in this project. Some tissues were processed for downstream analyses and some HMD regions were delegated to culture as patient-derived explants.

**Table 1.**
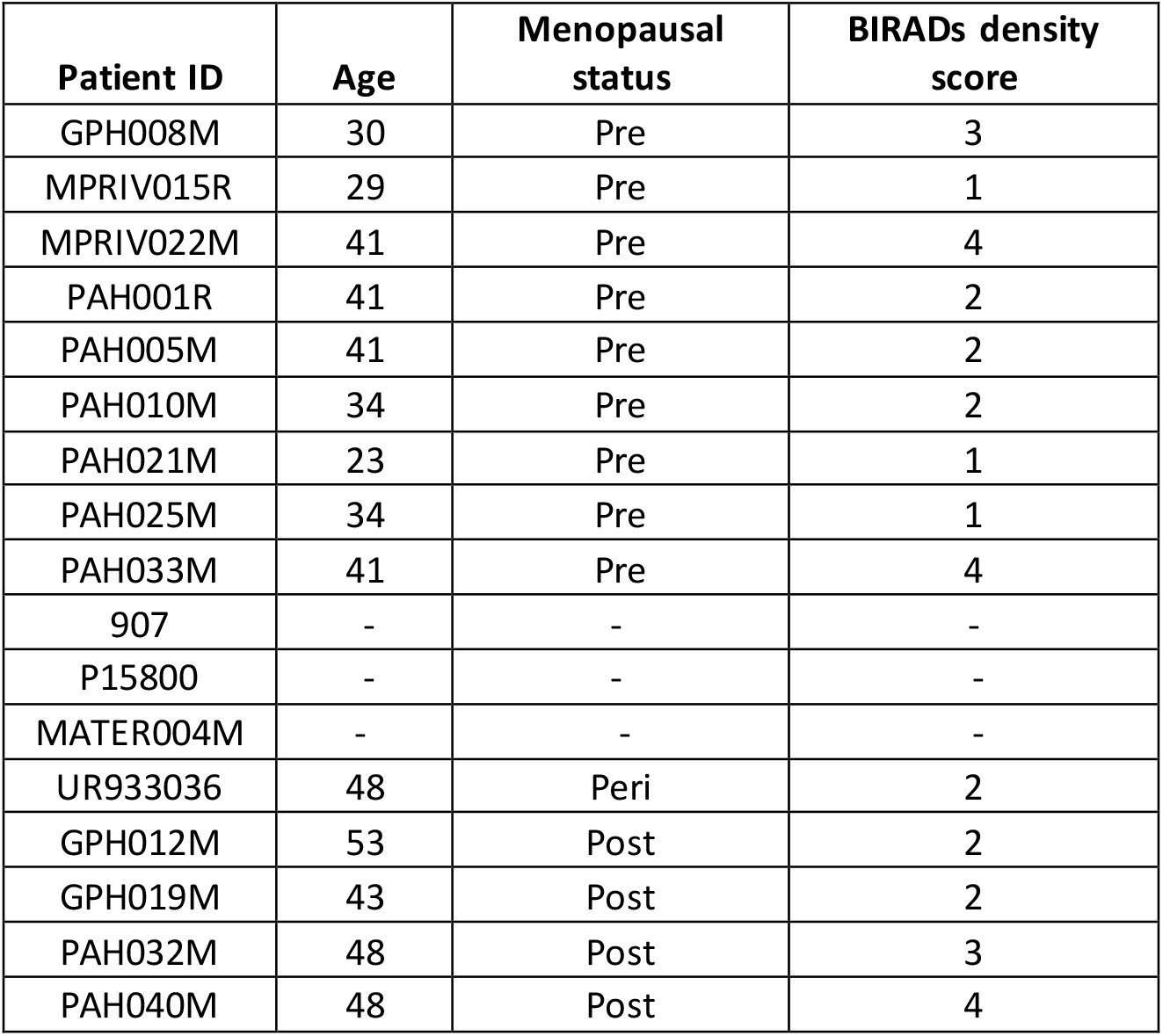
Patient demographics from which tissue was examined for this study.

| Patient ID | Age | Menopausal status | BIRADs density score |
| --- | --- | --- | --- |
| GPH008M | 30 | Pre | 3 |
| MPRIV015R | 29 | Pre | 1 |
| MPRIV022M | 41 | Pre | 4 |
| PAH001R | 41 | Pre | 2 |
| PAH005M | 41 | Pre | 2 |
| PAH010M | 34 | Pre | 2 |
| PAH021M | 23 | Pre | 1 |
| PAH025M | 34 | Pre | 1 |
| PAH033M | 41 | Pre | 4 |
| 907 | - | - | - |
| P15800 | - | - | - |
| MATER004M | - | - | - |
| UR933036 | 48 | Peri | 2 |
| GPH012M | 53 | Post | 2 |
| GPH019M | 43 | Post | 2 |
| PAH032M | 48 | Post | 3 |
| PAH040M | 48 | Post | 4 |

### 2.2 RNA extraction and RT-qPCR

RNA was extracted from human breast tissue after homogenization into TRIzol (Ambion, Life Technologies) using Qiagen Tissue Lyser II with metal beads. At the 1:1 isopropanol: sample step, the solution was added to a Bioline RNA extraction column (Catalog no BIO-52075) to purify RNA according to manufacturer’s instructions. RT-qPCR was performed as previously described (Hugo, Pereira et al. 2013). Primer sequences are detailed in Table 2.

**Table 2.**
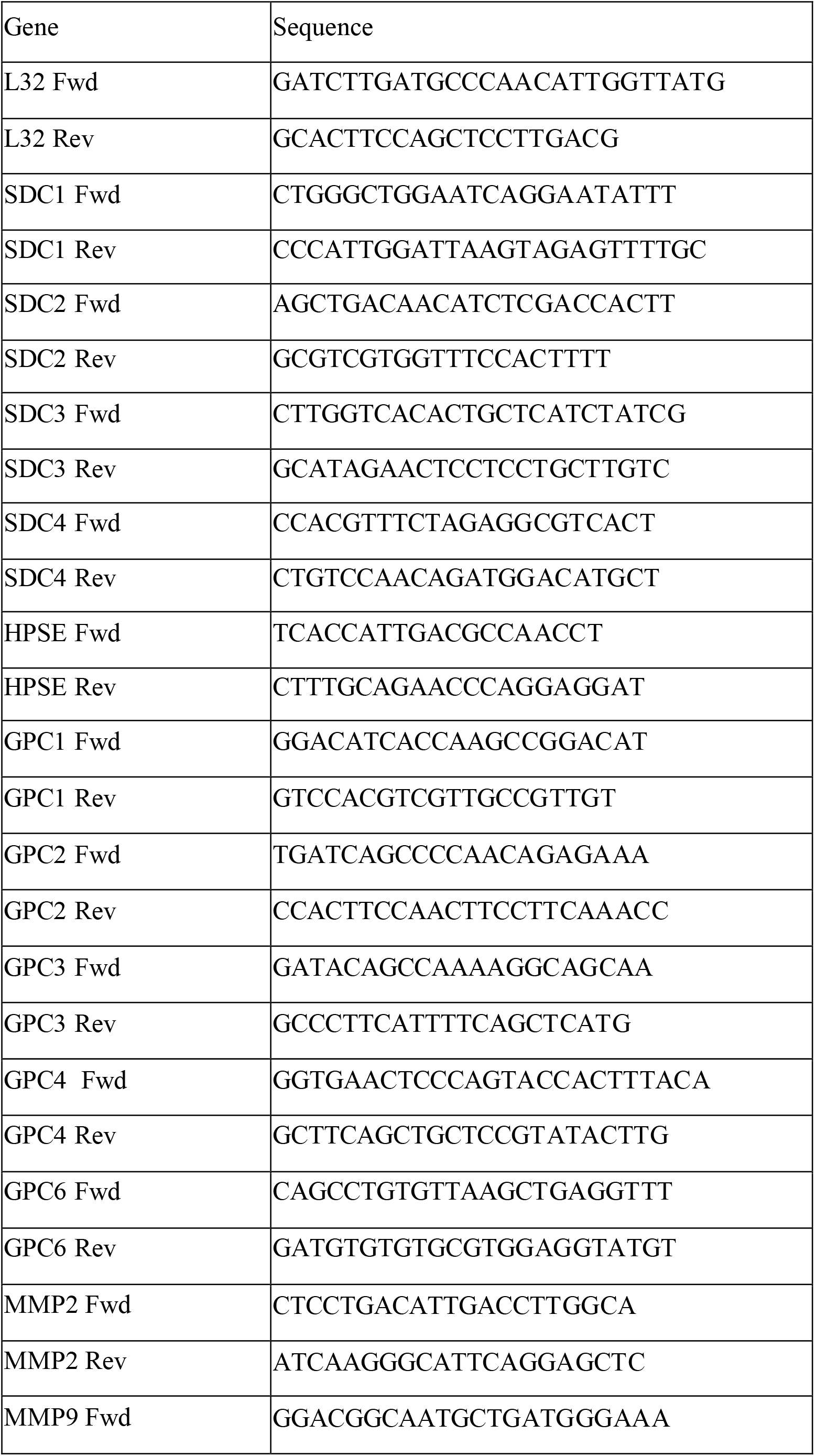
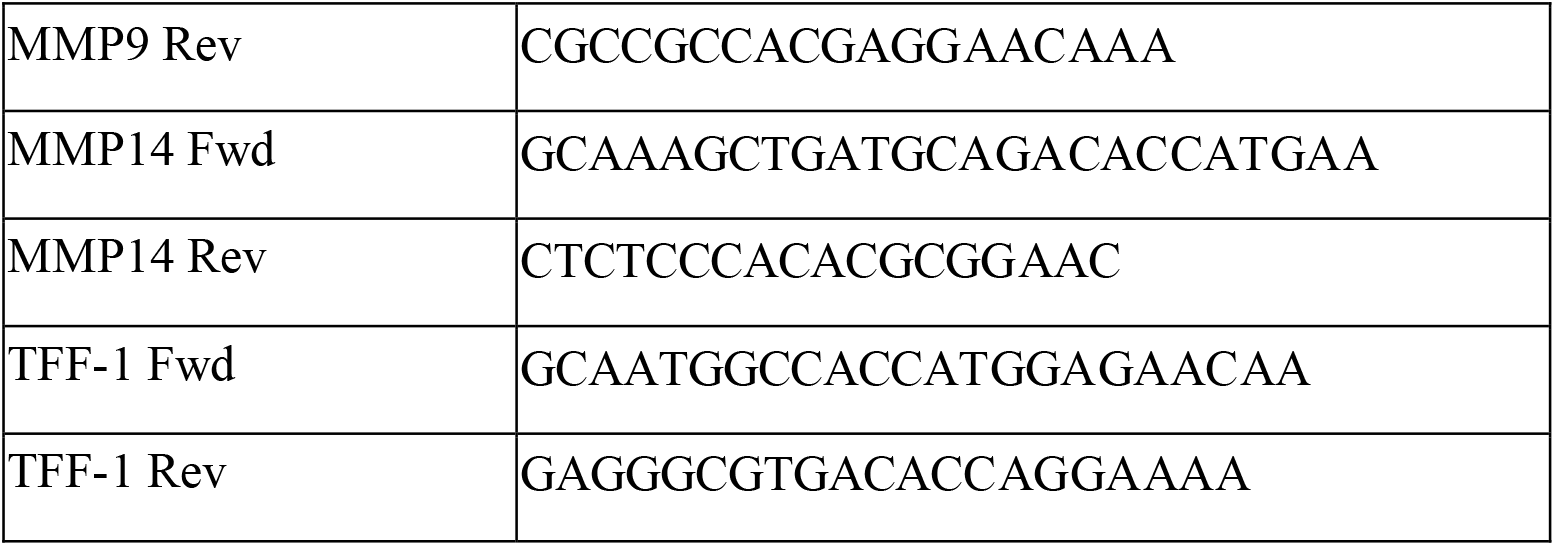
Sequences of RT-qPCR primers used in this study.

**Table 3.** Primary antibodies used in this study for Immunohistochemistry.

| **Antigen** | **Antibody** | **Dilution** | **Supplier** |
| --- | --- | --- | --- |
| SDC1 | Monoclonal Rabbit Anti-Human | 1:100 | Abcam (ab130405) |
| SDC4 | Polyclonal Rabbit Anti-Human | 1:100 | Abcam (ab24511) |
| TFF1 | Polyclonal Rabbit Anti-human | 1:1000 | Invitrogen (PA5-31863) |
| HPSE (Hpa) | Monoclonal Mouse Anti-Human | 1:100 | ThermoFisher Scientific (MA5-16130) |

### 2.3 Immunohistochemistry

Cut sections of HMD vs LMD, were fixed overnight in 4% paraformaldehyde made up in PBS at 4 degrees, processed to paraffin, sectioned and stained with various primary antibodies (detailed in Table 2) on an automated system (Ventana Discovery Ultra, Roche, Switzerland). “Intensity” (as specified on Y axes in Figure 1, parts A-E) represents per cent DAB positivity per tissue area, and was determined using ImageJ, where DAB brown positive tissue regions were separated using a Color Deconvolution plugin (H DAB), a threshold applied, and % positive area quantified.

**Figure 1.**
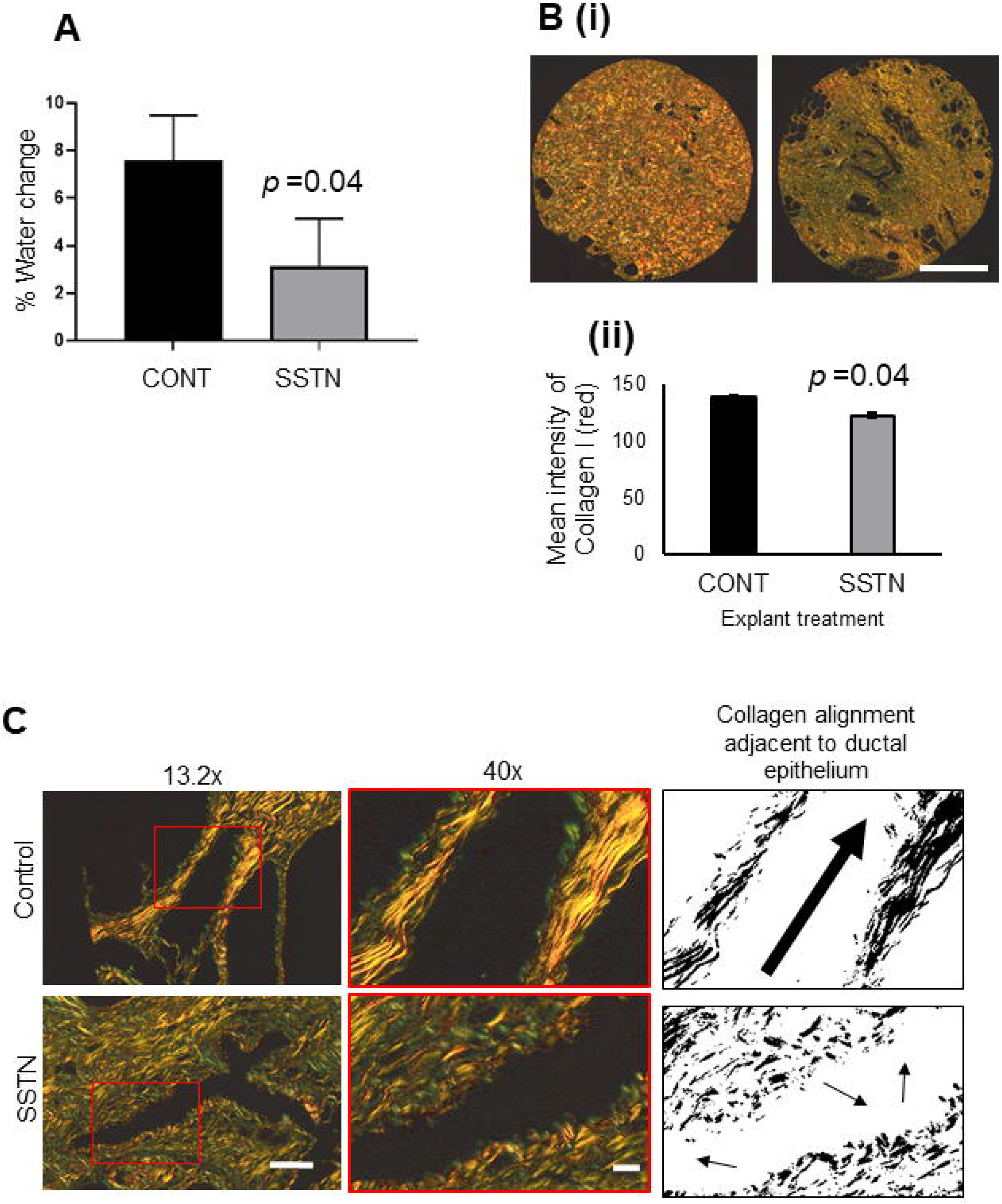
HPSG expression in HMD vs LMD. A. RT-qPCR for various HSPG proteins and HPSE (HPSE) in HMD versus LMD paired patient tissue. Fold change (HMD/LMD) gene expression from 15 patients, plotted on a logarithmic Y-axis. Delta CT values were obtained using L32 as the housekeeper gene, then converted to ddCT (2^CT^) for all fold change calculations. Student’s paired t test was used to determine significance, where ** is p<0.001 and *** is p<0.0001. Significance was achieved after adjustment for multiple testing. B. Assessment of SDC1 and SDC4 protein expression in HMD vs LMD tissues. Representative IHC images are displayed with intensity data (% DAB positivity per tissue area) plotted underneath. Lobular epithelium, n=4 pairs for SDC1, n=9 pairs for SDC4; C. Stromal regions, n=35 pairs for SDC1, n=30 for SDC4. D. Ductal vs lobular epithelium (HMD only shown) for SDC1 or SDC4, n=8 pairs for SDC1 and n=12 for SDC4. E. Ductal vs lobular epithelium (HMD only shown) for HPSE, n=12 pairs analysed. Student’s paired T test was used to determine significance, where p<0.001 is indicated by ** and p<0.0001 is indicated by ***. All images were captured at 10x magnification, scale bar = 50 μM.

### 2.4 Patient-derived explants of normal breast tissue

Briefly, ~2 cm^3^ sized pieces of breast tissue excised from high, medium or low MD regions, determined from a slice mammogram by a radiologist (TL), were cut into smaller pieces of approximately 0.5 cm^3^ in size and placed onto a gelatin sponge scaffold submerged in media inside a well of a 24-well culture plate. Viability of this tissue was then maintained for up to 2 weeks in 24 well plates, submerged in basal media as previously described (Centenera, Hickey et al. 2018) to sustain tissue viability and hormone responsiveness. For HPSE inhibitor treatment, this basal media comprised of RPMI containing 10% FBS (Thermofisher Scientific, Australia), 1 x penicillin and streptomycin (Merck, Australia), 100μg/mL hydrocortisone (Merck, Australia) and 100μg/mL insulin (Merck, Australia) was supplemented either one of the following: 100μM fondaparinux (sourced from Princess Alexandra Hospital Pharmacy), 100μM PI-88, 10μM PG545 control or 10μM PG545 (provided by V. Ferro). For estrogen (1nM Estradiol, Merck, Australia) and tamoxifen (1uM 4-hydroxy-tamoxifen, Merck, Australia) treatments, phenol red-free RPMI was supplemented with 10% charcoal-stripped FBS (Thermofisher Scientific, Australia), with all other supplements the same as for the HI experiments. For all treatments, media within 24 well plates was replenished twice weekly.

### 2.5 NMR measurements

The individual breast explants were removed from the underlying sponge and placed in a well within a dry 24 well plate for Portable NMR measurements. For PG545 and E2 treatments, NMR T1 values of the samples were measured using saturation-recovery pulse sequence on a PM5 NMR-MOUSE instrument (Magritek, Wellington, New Zealand) as previously described (Tourell, Ali et al. 2018, Huang, Ali et al. 2019). All saturation-recovery curves exhibited mono-exponential recovery. The curves were least-squares fitted with a three-parameter exponential fitting function, and the time constants of the recovery were taken as the respective T1 values. For SSTN treated explants, % water was measured using two approaches: (1) Carr-Purcell-Meiboom-Gill (CPMG) decay followed by an inverse Laplace transform, as described in (Ali, Tourell et al. 2019). This approach yielded T2 relaxation spectra with two peaks (Fat and Water), whose relative areas were taken as the content of the respective chemical component within the tissue; (2) Diffusion NMR measurements followed by two-component least-squares fitting of the diffusion attenuation plot (Huang, Ali et al. 2019). The amplitudes of the two components of the fit (Fat and Water) were taken as the content of the respective chemical component within the tissue.

### 2.6 Picrosirius red (PSR) staining and analysis under polarized light

PSR staining of human breast tissues to detect fibrillar collagen (collagen I) was performed as previously described (Kiraly, Hyttinen et al. 1997). Quantification of staining was performed using specific ImageJ macros written to identify and quantify red colour in PSR-stained sections illuminated with polarized light to visualize thick fibrillar collagens.

### 2.7 ELISA for shed SDC1 and SDC4 in explant media

SDC1 and SDC4 proteins shed into the conditioned media surrounding explanted normal mammary tissue at experimental endpoint (day 14) were quantified using the SDC1 and SDC4 ELISA kits (Raybiotech, USA) according to the manufacturer’s instructions, following dilution of the conditioned media 1:4 with the kit dilution buffer.

### 2.8 Statistics

Pearson’s r coefficients and respective p values and the Student’s T test (used to assess the differences in MD changes and gene expression levels between the different treatment groups) were calculated using GraphPad Prism 8.

## 3 Results

### 3.1 HMD vs LMD pairwise ECM gene and protein expression analysis

#### 3.1.1. RT-qPCR data

As shown in Figure 1, RT-qPCR assessment of a range of HSPGs showed that mRNA levels of SDC1 and SDC4 were significantly more abundant, 14-fold and 7-fold higher, respectively, in HMD versus LMD paired tissue from 15 women. In contrast, levels of GPC3 and GPC6 were significantly decreased by 0.8- and 0.65-fold, respectively. Significance was achieved after adjustment for multiple testing. The proteins coded by these mRNAs are substrates for HPSE. We did not find any difference in HPSE expression between HMD and LMD. Given that increased stroma is characteristic of HMD (Huo, Chew et al. 2015), postulated in this study to be driven by HSPG abundance, SDC1 and SDC4 were thus taken forward for further analysis.

#### 3.1.2. Localization of SDC1 and SDC4 in normal mammary tissue

Stromal, not epithelial, localisation of SDC1 protein has been implicated in promoting breast density and breast cancer (Lundstrom, Sahlin et al. 2006). Therefore, we performed a detailed IHC investigation of SDC1 and SDC4 PGs in epithelial and stromal compartments of paired HMD vs LMD tissues (Figure 1 parts B-E). No significant difference was observed for epithelial SDC1 or SDC4 protein expression in HMD versus LMD (Figure 1B), however SDC1 expression was higher in stromal regions derived from HMD compared with LMD (Figure 1C). This was not observed for SDC4, suggesting SDC1 abundance may drive HMD. Both SDC1 and SDC4 protein levels appeared more abundant in ductal rather than in lobular epithelia, irrespective of MD status (Figure 1D). No difference in HMD vs LMD epithelial versus stromal regions were observed for HPSE *(data not shown),* however HPSE displayed the same expression pattern as for SDC1 and SDC4 in that it was higher in ductal versus lobular epithelium (Figure 1E).

### 3.2 Establishment of a patient-derived explant model to assess MD change

All published reports that have examined MD change have been longitudinal and in human population cohorts (Junkermann, von Holst et al. 2005, Cuzick, Warwick et al. 2011, Jacobsen, Lynge et al. 2017). We sought a means to test the effect of exogenous agents on MD in a culture setting using intact mammary tissue, and adopted a whole-tissue explant model (patient-derived explant, PDE), originally designed for the study of human cancers *ex vivo* (Centenera, Hickey et al. 2018). These original studies describe the culture of cancer tissue for periods of time measured in days, however we hypothesized that to observe any measurable change in density in normal tissue, treatment times would need to be extended to weeks. Initial concern in adopting this extended treatment time was that tissue viability would be compromised. However, as shown in Figure 2A (i), morphology of the PDEs after 2 weeks was comparable to tissue at day 0, and we found minimal activated caspase-3 activity in medium MD explant tissue cultured for 2 weeks, indicating that apoptosis was not a major event in these tissues during this time (Figure 2A, ii).

**Figure 2.**
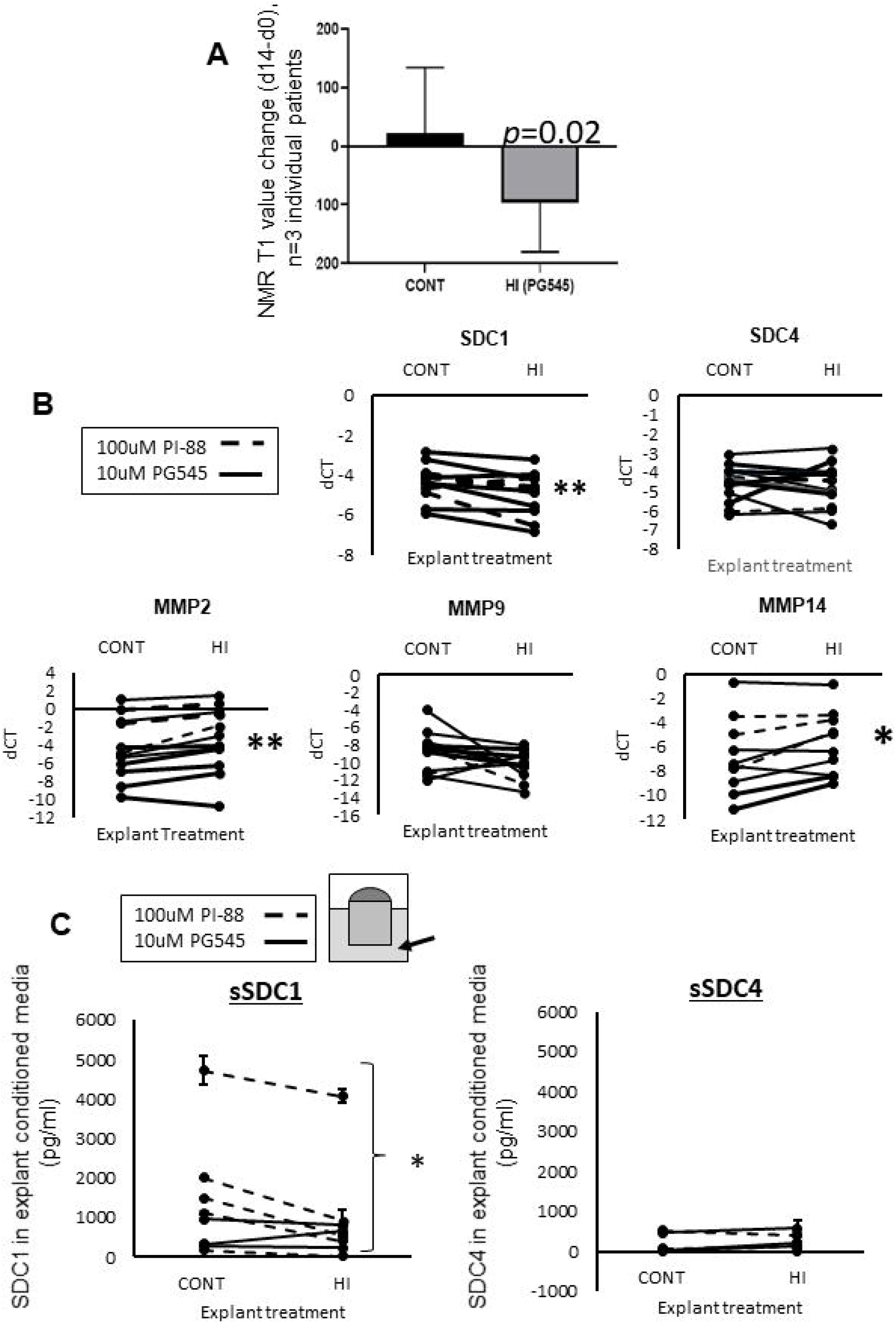
A. (i) Morphological comparison of original tissue at t=0 and following 14 days as explant culture. S denotes position of the sponge, delineated from the explant tissue by a dotted line. (ii) Viability of explanted tissue up to 14 days as assessed by % activated caspase 3 activity as measured by IHC. C= clear media (Phenol-red free); r= red media (Phenol-red containing). B. 1nM Estradiol treatment of PDEs led to a clear upregulation of the E2-signaling pathway protein TFF-1, images and quantification are typical representative data of 3 individual patients. C. Mammographic density change (d14-d0) in E2 treated mammary tissue explants as measured by the change in NMR T1 values. For part C only, Student’s paired T test was used to determine significance, with p values displayed on graph, n=3 individual patient tissue tested. For all images shown, 10x magnification was used, scale bar = 50 μM.

#### 3.2.1 Increase in MD with estradiol treatment of PDEs

We then treated our medium MD PDEs with 1nM estradiol for 14 days, as estrogen is an implicated driver of MD (Greendale, Reboussin et al. 1999, Byrne, Ursin et al. 2017). After 14 days, we found the estrogen receptor signaling pathway mediator TFF-1 protein to be upregulated in comparison to the control (Figure 2B).

We have demonstrated that NMR T1 (and to a lesser extent, T2) values correlate strongly with microCT-derived % HMD in ex vivo explants, such that T1 values may be used as a surrogate marker for MD (Huang, Ali et al. 2019). We therefore chose to measure T1 as a surrogate marker of MD in our PDE model at t=0 and t=14, to determine MD change. As shown in Figure 2C, in all three experiments in which tissue from women was tested, estrogen increased MD change (d14-d0) compared to control treated (p=0.04). In another patient, tamoxifen negated estradiol-mediated MD increases over the experimental period (14 days, Supplementary Figure 1).

#### 3.2.2. Coupling of HPSE with SDC1 and SDC4 induction with ERα pathway activation

Since it is known that estrogen upregulates HPSE mRNA (Elkin, Cohen et al. 2003), that HPSE is a key modulator of both SDC1 and SDC4 abundance in the breast (reviewed in (Sarrazin, Lamanna et al. 2011), and that cleavage of SDC1 leads to SDC1 mRNA induction (Ramani, Pruett et al. 2012), we investigated whether the E2-HPSE-SDC1 axis could be relevant in our human mammary tissue explants. As shown in Figure 3A, significant correlations were observed for both SDC1 and SDC4 with HPSE in E2-treated explants. Each of the data points shown in this Figure represent the average of 3 explants from 1 patient (independent biological replicates), and in Figure 3A, data from 8 individual patients was analysed. The correlation observed in Figure 3A was lost when tissue explants were either treated with tamoxifen alone (Figure 3B) or co-treated with E2 and tamoxifen (Figure 3C), where 6 individual patient’s explant tissue was examined. Furthermore, HPSE protein was found to be more abundant in E2-treated PDEs (supplementary Figure 2A). These findings suggest a coupling of HPSE with SDC1 and SDC4 induction where the ERα pathway was activated, however, further studies with larger numbers are required to confirm this, including a more thorough examination of *in vitro* and *in vivo* tissues.

**Figure 3.**
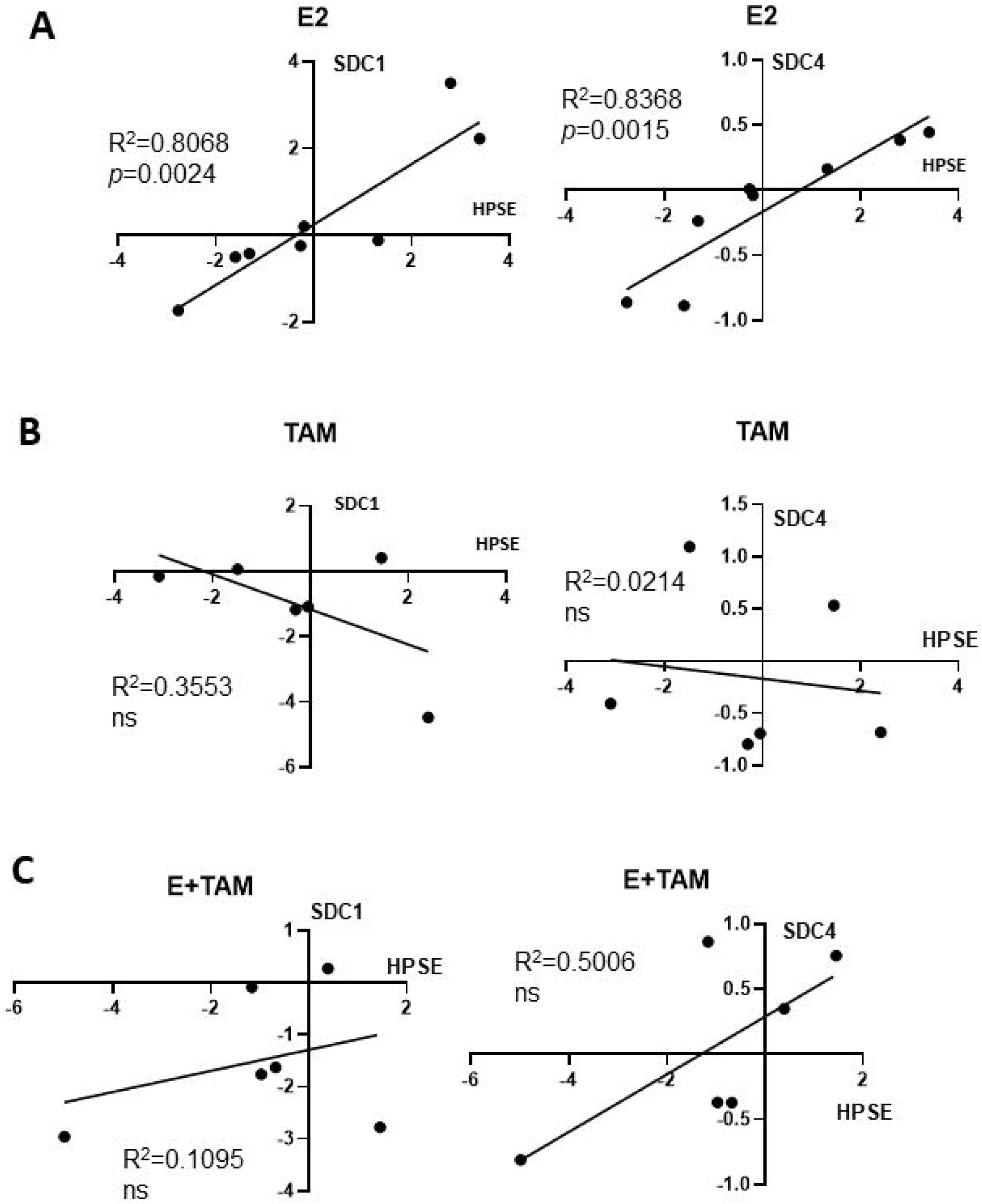
RT-qPCR data from hormone treated explants: Correlation of HPSE with SDC1 or SDC4 in E2 (A), Tamoxifen (B) and E2+Tamoxifen (C) treated explants. Expression data shown is normalized to control-treated explant mRNA control (ddCT). Each data point represents the average expression from 3 explants from 1 patient, where n=8 for part A, n=6 for parts B and C. Pearson’s r coefficient is shown and where p values stated where p<0.05; ns = not significant.

### 3.3 Effect of HPSE inhibition in PDEs

Given that we were able to detect an increase in MD in the PDE model with estradiol, and that HPSE regulation of SDC may be implicated in mediating this change, we decided to use the PDE model to inhibit HPSE specifically and assess the effect on MD and gene expression outputs.

#### 3.3.1 HPSE inhibition reduced MD

We examined the effect of the HS mimetic PG545 (Chhabra and Ferro 2020) in the PDEs for the same period of time as for estradiol (14 days). IHC assessment of HPSE revealed that it was predominantly located around glandular epithelia, and 10uM PG545 led to the strongest proportional reduction in peri-glandular HPSE protein content (Supplementary Figure 1A). Therefore, 10uM was used in further studies. As shown in Figure 4A, in all three experiments in which breast tissue from women was tested, PG545 decreased MD relative to the respective control, as indicated by NMR T1 change (d14-d0, p=0.02).

**Figure 4.**
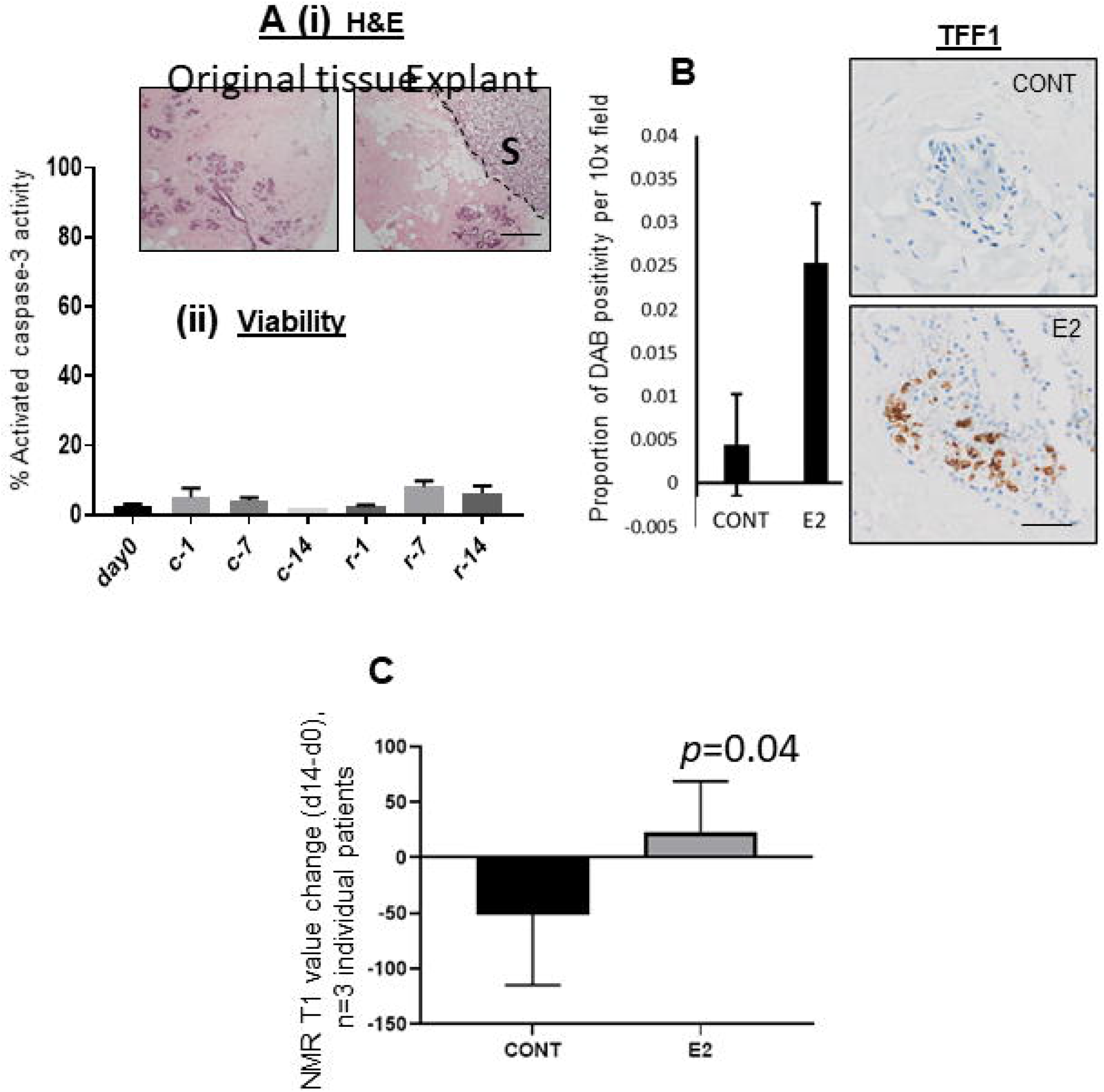
A. MD change (d14-d0) in PG545 treated mammary tissue explants as measured by the change in NMR T1 values, Student’s paired T test was used to determine significance, with p values displayed on graph, n=3 individual patient tissue tested. B. RT-qPCR data from HI treated explants. Dotted line denotes 100uM PI-88 treated explants, where control is 100uM Fondaparinux; solid line denotes 10uM PG545 treatment, where control is media alone. More gene expression data is found in supplementary Figure 3. Delta CT values are plotted, determined using using L32 as the housekeeper gene. C. Inhibition of HPSE led to a decrease in the abundance of shed SDC1 in explant conditioned media (Student’s paired T test used to determine significance, * denotes p<0.05). Dotted line: PI-88, solid line: PG545. For SDC1, n=8 pairs, for SDC4, n=6.

An additional HS mimetic (PI-88) was also used in PDEs (separate to PG545). PI-88 has a similar mechanism of action (Chhabra and Ferro 2020) and the synthetic pentasaccharide fondaparinux was used as a control for this drug. Efficacy in reducing HPSE protein level by IHC is shown in Supplementary Figure 2B.

#### 3.3.2 Gene expression changes following HPSE inhibitor treatment in PDEs

Analysis of the effects of HPSE inhibition on various HSPG mRNA levels in explanted tissue at endpoint (14d) by RT-qPCR revealed a significant reduction in SDC1 expression, but not in SDC4 or other HSPGs (Figure 4B, Supplementary Figure 3), again indicating a complex transcriptional interplay between HSPG and SDC1. Given that HPSE activity has also been shown to positively influence MMP14 expression (Gomes, Bhat et al. 2015) and is associated with MMP9 (Abu El-Asrar, Siddiquei et al. 2016) we also examined MMP expression in the control versus treated tissues. Of the MMPs that displayed expression above baseline (MMP-2, −9 and −14), MMP2 and MMP14 showed a significant increase after HPSE inhibitor (HI) treatment (Figure 4B).

#### 3.3.3. Effect of HPSE inhibitor treatment on shed SDC1 and SDC4

The abundance of SDC1 or SDC4 protein was examined in the conditioned media of PDEs at t=14, as an indicator of HPSE activity, given that the form of the protein detected by this means has been shed from the cell membrane (Yang, Macleod et al. 2007).

As shown in Figure 4C, there was, on average, 2 to 8-fold more SDC1 than SDC4 protein shed into the media, and only SDC1 displayed a significant reduction in abundance with HPSE inhibition (p<0.05).

### 3.4. SDC1-specific inhibition using SSTN reduced fibrillar collagen and MD

Given that gene expression and shedding for SDC1 (but not SDC4) was significantly reduced with heparanase inhibition, we hypothesized that SDC1 was mediating the observed MD change shown in Figure 4A. But how could SDC1 mediate MD? McConnell and colleagues found that peri-ductally aligned collagen is correlated with MD (McConnell, O’Connell et al. 2016), and a direct role for membrane-bound SDC1 in physically aligning collagen fibres has been described (Yang and Friedl 2016). We therefore decided to interrogate the connection between SDC1 and collagen and MD in our PDE model.

SSTN is a peptide designed and validated to block SDC1-binding to ITGαVβ3, and hence ITGαVβ3 binding to peri-ductal collagen, but also with Insulin-like growth factor 1 receptor (IGF1R) (Rapraeger 2013). First, we validated its effectiveness in our own hands to block MDA-MB-231 breast cancer cell binding to vitronectin, an interaction that specifically requires SDC1-ITGαVβ3 complexes. As shown in Supplementary Figure 4, 30uM of SSTN reduced MDA-MB-231 cell binding and this concentration was therefore carried forward into our PDE experiments.

As shown in Figure 5A, SSTN significantly decreased MD relative to the respective control, as indicated by % water change determined by NMR (d14-d0, n=3 individual patients, p=0.02). PSR staining enhances the natural birefringence of collagen bundles. When visualized under polarized light, PSR-stained collagen I appears red while collagen III appears green, and thus PSR staining can be used to determine collagen I/III content (Vogel, Siebert et al. 2015). SSTN led to a visible reduction in dense collagen, or fibrillar collagen content, as measured by PSR visualized under polarized light (Figure 5B, parts i and ii). Upon closer inspection, the arrangement of collagen adjacent to ductal epithelium in SSTN treated explants exhibited reduced alignment compared to the control (Figure 5C).

**Figure 5.**
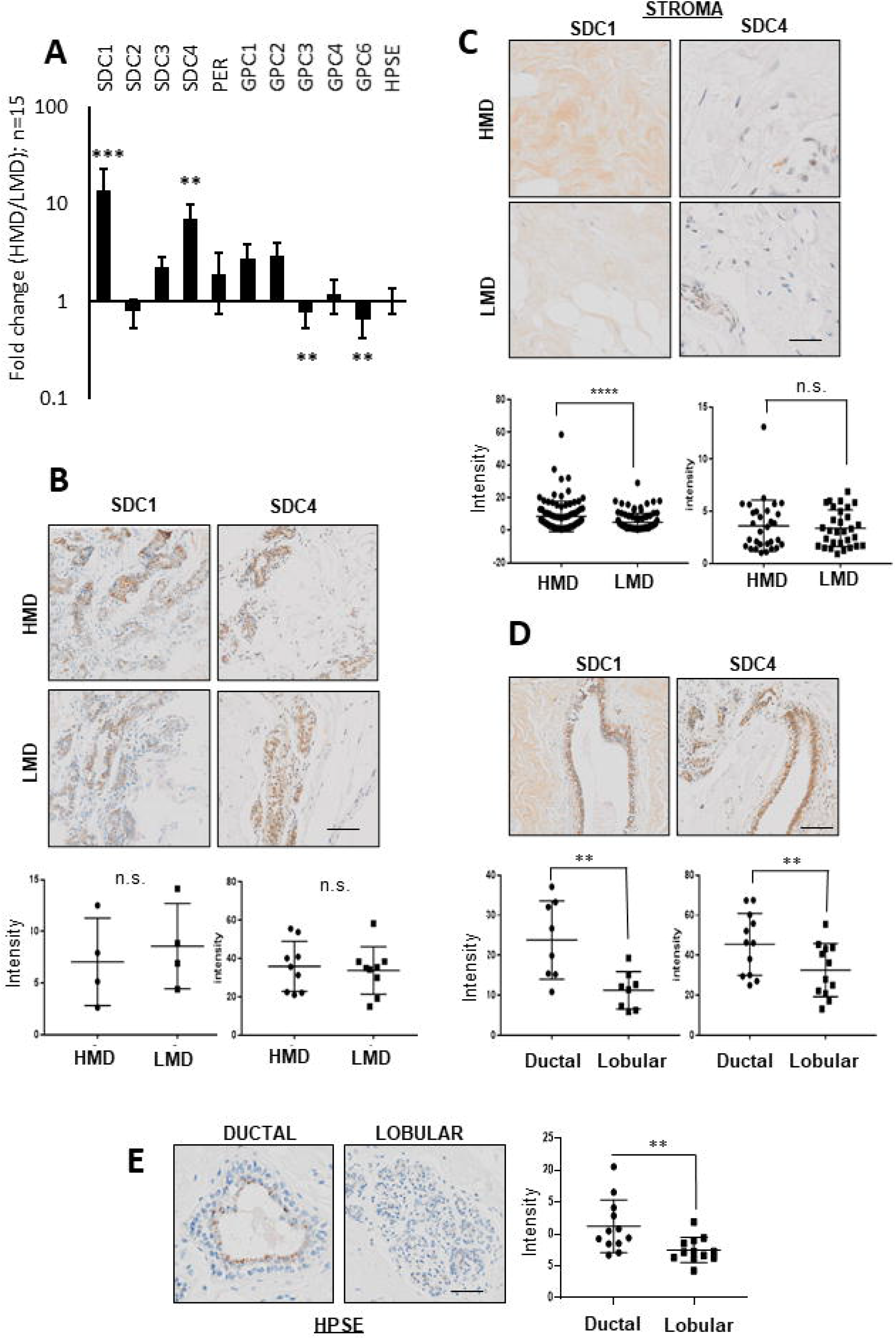
A. MD change (d14-d0) in SSTN treated mammary tissue explants as measured by the change in % water diffusion values as determined by single-sided NMR. B. (i) Picrosirius red staining of PDE tissue microarray and imaging with polarized light to visualize dense collagen fibres; (ii) quantification of dense collagen represented as the mean intensity of red fibres visualized by polarized light (data from n=3 patients, 3 explants quantified per patient). Results are the mean and error bars represent standard error. P value was determined using Student’s paired T test. Images captured at 4x magnification, scale bar = 200 μm. C. Picrosirius red images, enlarged region of image on left is within red box, then black and white thresholded image to highlight changes to alignment of collagen adjacent to ductal epithelium after treatment with Synstatin. Images captured at 13.2 x magnification, scale bar = 100 μM, 40 x magnification, scale bar = 20 μM.

## 4 Discussion

We show for the first time that MD change can be measured *ex vivo.* This significant finding has enabled a dissection of molecules involved in the maintenance of MD. Our studies illustrate a key role for HPSE in maintaining MD, with SDC1 and possibly SDC4 implicated in this process. We have previously shown that stroma is the key differentiator between high versus low MD (Huo, Chew et al. 2015), and we show in the current study that SDC1 and −4 are both more abundant in HMD vs LMD (Figure 1A), and that SDC1, but not SDC-4, was more abundant in the HMD stroma (Figure 1C). Although the E2-HPSE-SDC induction pathway induced SDC1 and 4 in a relatively equal manner (Figure 3A), in HPSE inhibitor-treated explant tissue, SDC1, but not SDC4, gene expression was significantly reduced (Figure 4B), and shedding of SDC1, but not SDC4, was reduced (Figure 4C). Finally, our results indicate that fibrillar collagen is important in the maintenance of MD, and that SDC1 plays a direct role in mediating this effect, as SSTN reduced MD and fibrillar collagen abundance (Figure 5).

Despite both SDC1 and SDC4 mRNA being more abundant in HMD vs LMD (Figure 1A), HPSE modulation had a more significant, and hence functional effect on SDC1 rather than SDC4. Interestingly, SDC1 (not SDC4) is implicated in both MD and BC (reviewed in (Shawky, Ricciardelli et al. 2015)) where SDC1 is informative as to BC staging (Cui, Jing et al. 2017). Our work highlights that SDC1 may be a key molecular target in efforts to reduce mammographic density as a means to reduce the associated BC risk.

Shed SDC1 has a number of potential effects in promoting tumourigenesis: growth factors bound to the extracellular domain can be carried into the nucleus and potentiate proliferation (Stewart, Ramani et al. 2015) and also increase wnt signaling (Alexander, Reichsman et al. 2000). HPSE induces SDC4 shedding, as we observed in the conditioned medium of our PDEs (Figure 4C), however the pathological significance of SDC4 shedding is in cardiac disease (Strand, Aronsen et al. 2015). Given that shed SDC1 in the serum is a prognostic factor in several cancers (Joensuu, Anttonen et al. 2002, Vassilakopoulos, Kyrtsonis et al. 2005, Szarvas, Reis et al. 2016) it is likely that there are several other pro-tumourigenic effects of shed SDC1 into the microenvironment that are yet to be discovered. SDC1 shedding itself increases SDC1 expression in the cell (Ramani, Pruett et al. 2012), but may also increase SDC1 expression in neighbouring cells, such as fibroblasts in the stroma, a mechanism through which stromal SDC1, particularly in HMD, may be potentiated, as we have observed (Figure 1C).

We found that SDC1, SDC4 and HPSE were significantly more abundant in ductal epithelium compared with lobular epithelium (Figure 1D, E). The basis of this expression pattern is not clear, however, it may be related to differing mechanical pressure placed on ductal versus lobular epithelia. McConnell and colleagues (McConnell, O’Connell et al. 2016) found significant correlations between MD and localised PSR enhanced collagen birefringence located around breast ducts (R2=0.86, p<0.0001), but not around lobules or within distal stromal regions (R2=0.1, p=0.33; R2=0.24 p=0.14, respectively). The presence of this restrictive collagen lining may create a greater force on the cells underlying it. In the same way that increased load in cartilage due to exercise leads to an increase in proteoglycan content (Bird, Platt et al. 2000), perhaps this force is responsible for the observed SDC1 and SDC4 expression patterns. Indeed, we see the expression of all syndecan family members increase in human mammary epithelial cells cultured on stiff versus soft 2D surfaces (6kPa vs 400Pa, *unpublished observations*).

We observed an increase in MMP2 and MMP14 in HPSE inhibitor treated explants (Figure 4B). Similarly, HPSE knockout mice have been reported to display an upregulation of MMP2 and MMP14, where these metalloproteinases are thought to compensate for the loss of HPSE (Zcharia, Jia et al. 2009). Although decreased, SDC1 shedding was not eliminated in HPSE inhibitor-treated explants (Figure 4C). In light of the Zcharia study (Zcharia, Jia et al. 2009), this suggests that some SDC1 shedding was possible in the HPSE inhibitor-treated explants due to a MMP2 and −14 compensatory mechanism.

This study has demonstrated that the HPSE inhibitors PG545 and PI-88 are useful to interrogate the mechanistic effects of HPSE in tissue samples on MD. However, these inhibitors have undesirable anti-angiogenic and other off-target effects when used systemically (Chhabra and Ferro 2020) and thus are unlikely MD-reducing agents to take forward to clinical trials. Tamoxifen reduces MD (Shawky, Martin et al. 2017), however, its long-term use is associated with a wide range of potential side effects, some of which are intolerable (Thorneloe, Hall et al. 2020). Both of these approaches are therefore potentially not feasible for MD reduction, although their mechanism of action provides important insight into how MD is governed, as our work alludes to, via the action of HPSE. We have uncovered a more targeted approach to MD reduction, discovered by teasing apart the connection between SDC1 and collagen. We believe this work will pave the way forward for other kinds of targeted approaches to be developed with aim to reduce MD and thus breast cancer risk.

## 6 Contribution to the field statement

Mammographic density (MD), also called breast density, describes the white areas on a mammogram. The amount of MD can reduce the ability of a radiologist to detect a breast cancer (BC), but also can make BC grow more aggressively. High MD is characterized by a preponderance of fibrous stromal tissue within the breast, which is rich in molecules called heparan sulfate proteoglycans (HSPGs). The abundance of these molecules is governed by the activity of the enzyme heparanase (HPSE). Estrogen is known to drive MD, and also to drive HPSE abundance. We show for the first time that an *increase* in MD can be detected *ex vivo*, using NMR technology in cultured human mammary tissue treated with estrogen. We show that inhibitors of HPSE, which sits downstream of estrogen, has the opposite effect. We show that the heparan sulfate proteoglycan (HSPG) syndecan-1 is the key protein that mediates the effect of HPSE modulation on MD. Taken together, this study supports a role for HPSE and SDC1 as novel mediators of MD. Molecular targeting of HPSE and/or SDC1 may be of use in reducing MD in women and thus improving BC detection via screening mammography but also restraining BC progression.

## 7 Author Contributions

HJH and EWT equally conceived the study design and managed its progress. XH, GR and HJH obtained human breast tissue from respective Pathology Departments of the Princess Alexandra Hospital and Mater Private Hospital and set up human breast tissue explants for two week culture. XH and HJH and maintained explant cultures, performed gene expression and together with GR, performed IHC intensity quantification. HJH performed ELISA for SDC1 and SDC4. XH and KIM obtained NMR T1 measurements and KIM performed analyses on this data. CES provided advice on estradiol concentrations and signaling outputs. RKO performed RT-qPCR data for Figure 1A. TB provided input as to the correct statistical method to use. TL assessed breast tissue slice mammograms, and determined valid areas of HMD vs LMD for excision. WDT and TEH provided hormonal reagents and technical advice on explant culture. LH provided HSPG primers and antibodies and advice in IHC interpretation. VF provided HPSE inhibitors used in explant culture, and advice in concentration optimisation. All authors reviewed the manuscript prior to submission.

## 9 Acknowledgments

The authors wish to thank the Translational Research Institute Histology Core for performing IHC contained in this manuscript, but also Dr Jason Northey for sharing the code he wrote for ImageJ to enable quantification of picrosirius red images for collagen I content. We thank Ms Gillian Jagger for her central role in recruiting, obtaining consent and co-ordinating the collection of breast tissue from the study participants. Finally, we thank the ladies who donated their tissue that enabled this study be performed.

---

**Supplementary Figure 1.** A. Dilution series (0- 20uM) of the heparan sulfate mimetic (heparanase inhibitor) PG545 and its effect on HPSE protein abundance, after 14 days treatment, as determined by IHC. Magnification 10x, scale bar = 50 μM. B. % water diffusion change data from 1 patient showing that the combination of tamoxifen with estrogen ameliorated MD increase over the treatment period (14 days).

**Supplementary Figure 2.** HPSE protein abundance as measured by IHC in A. PI-88 (HPSE inhibitor) and B. E2 treated explants. % DAB positivity was quantified from at least 5 10x microscope fields. Magnification 10x, scale bar = 50 μM.

**Supplementary Figure 3.** A. RT-qPCR data derived from HI treated explants for the various other HSPG family members where no change in expression was observed.

**Supplementary Figure 4.** MDA MB 231 adherence to two-dimensional, Vitronectin coated substrates in the presence of increasing concentration of the SDC1 inhibitor SSTN. A. Cellular morphology and B. % adherence calculations from images shown in A. Magnification 20x, scale bar = 50 μM.

