## Supplementary figures and images for "Heparanase promotes Syndecan-1 expression to mediate fibrillar collagen and mammographic density in human breast tissue cultured *ex vivo*"

### Supplementary figure 1

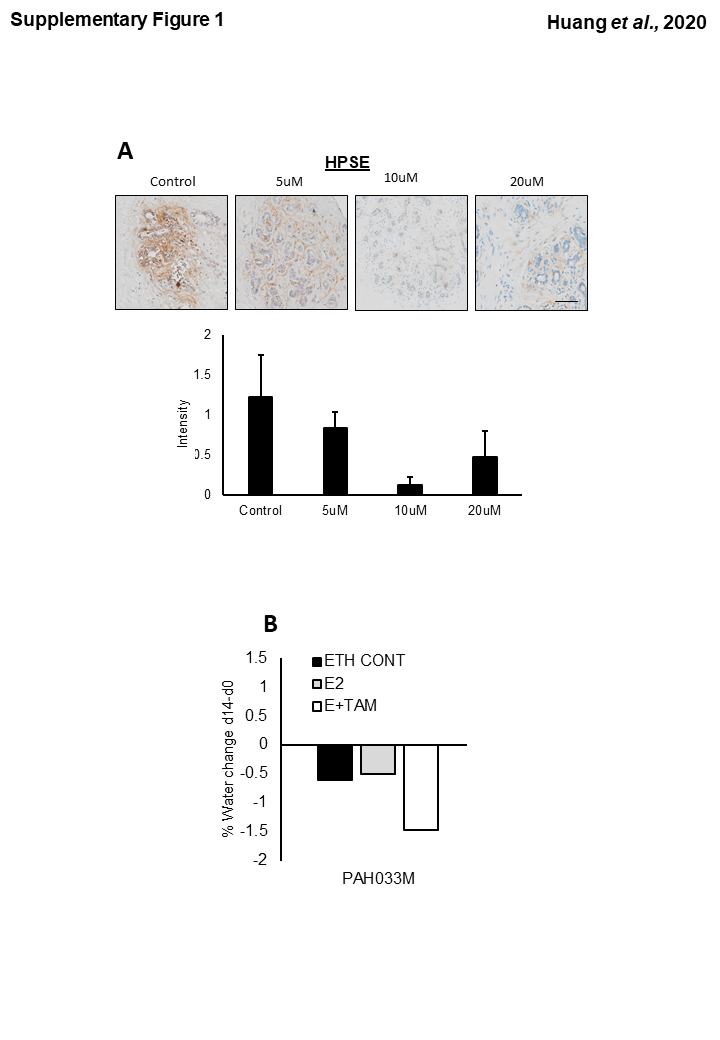

### Supplementary figure 2

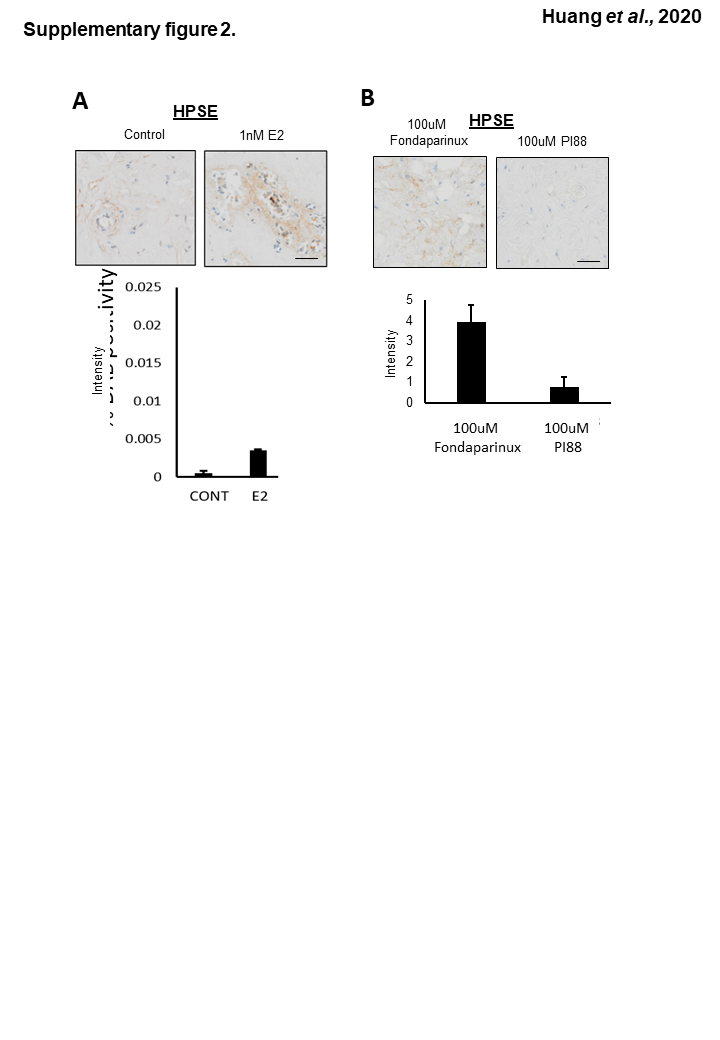

### Supplementary figure 3

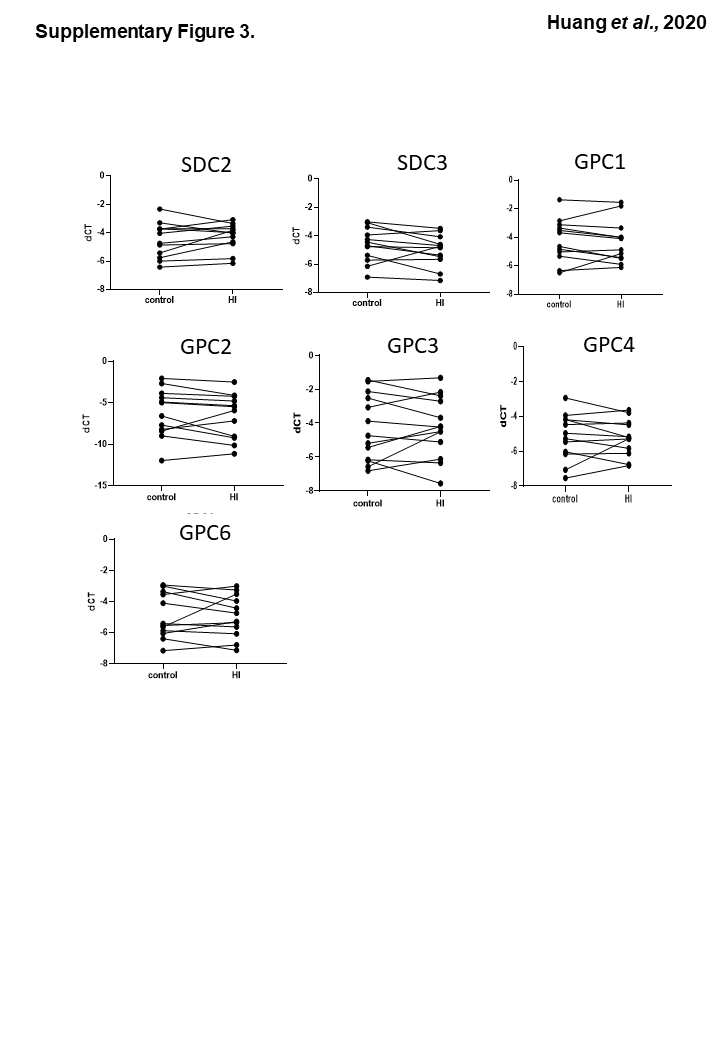

### Supplementary figure 4

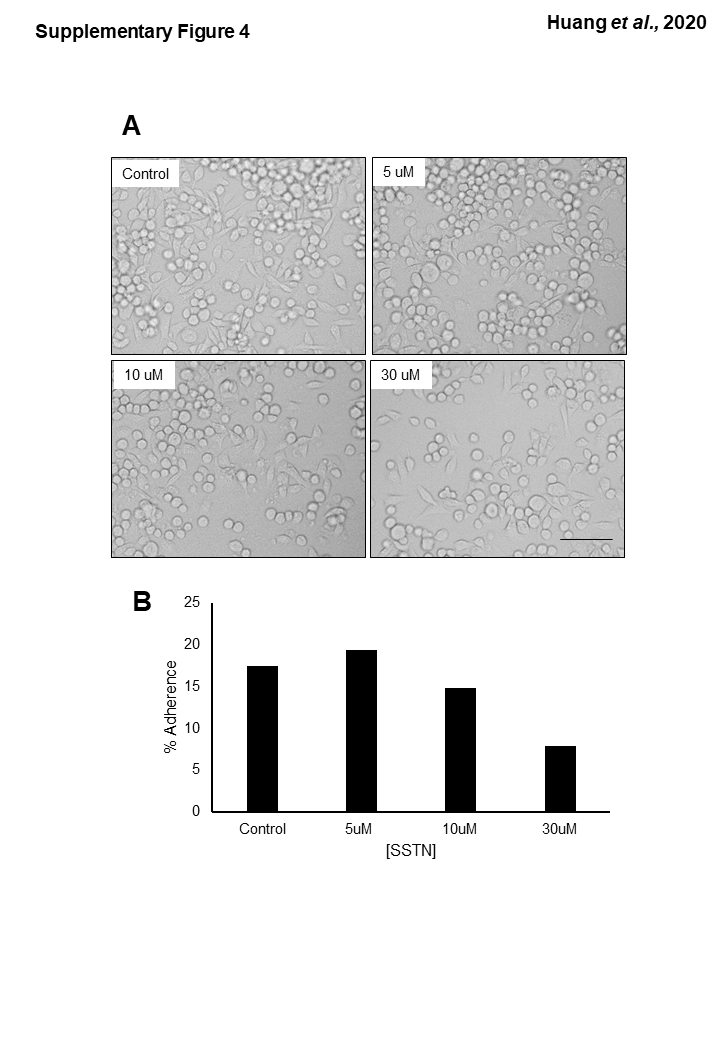
